# Therapeutic immunization with a whole cell vaccine reduces pneumococcal nasopharyngeal density, shedding, and middle ear infection in mice

**DOI:** 10.1101/2023.04.05.534641

**Authors:** Jayne Manning, Sam Manna, Eileen M Dunne, Viktoria Bongcaron, Casey L Pell, Natalie L Patterson, Sacha D Kuil, Poshmaal Dhar, David Goldblatt, E Kim Mulholland, Paul V Licciardi, Roy M Robins-Browne, Richard Malley, Odilia Wijburg, Catherine Satzke

## Abstract

Pneumococcal Conjugate Vaccines (PCVs) have substantially reduced the burden of disease caused by *Streptococcus pneumoniae* (the pneumococcus). However, protection is limited to vaccine serotypes, and when administered to children who are colonized with pneumococci at the time of vaccination, immune responses to the vaccine are blunted. Here, we investigate the potential of a killed whole cell pneumococcal vaccine (WCV) to reduce existing pneumococcal carriage and mucosal disease when given therapeutically to infant mice colonized with pneumococci. We show that a single dose of WCV reduced pneumococcal carriage density in an antibody-dependent manner. Therapeutic vaccination induced robust immune responses to pneumococcal surface antigens CbpA, PspA (family 1) and PiaA. In a co-infection model of otitis media, a single dose of WCV reduced pneumococcal middle ear infection. Lastly, in a two-dose model, therapeutic administration of WCV reduced nasal shedding of pneumococci. Taken together, our data demonstrate that WCV administered in colonized mice reduced pneumococcal density in the nasopharynx and the middle ear, and decreased shedding. A vaccine with similar properties in children would be beneficial in low and middle-income settings where pneumococcal carriage is high.

**Importance:** Although typically asymptomatic, pneumococcal carriage plays an essential role in transmission and the development of disease. Pneumococcal Conjugate Vaccines (PCVs) have reduced the burden of pneumococcal disease worldwide. However, their use has increased carriage and disease caused by non-vaccine serotypes, prompting investigations into serotype-independent pneumococcal vaccines. An additional limitation of PCVs is immune hypo-responsiveness to vaccines in children carrying pneumococci at the time of vaccination. Therefore, there is great interest in next generation vaccines such as whole cell vaccines. In this study we investigate a pneumococcal whole cell vaccine (WCV) for it effect on carriage in mice that are already colonized at the time of vaccination. We show that this ‘therapeutic’ vaccination of mice can reduce pneumococcal carriage density, shedding and infection of the middle ear. Our study suggests that WCV could be beneficial in high burden settings where carriage at the time of vaccination is more common.

## Introduction

*Streptococcus pneumoniae* (the pneumococcus) is a leading cause of morbidity and mortality worldwide (1). Pneumococci colonize the nasopharynx (carriage), from where they can disseminate to other tissues and cause diseases including otitis media and pneumonia (2). Compared with children from higher income settings, children in low and middle-income settings are colonized by pneumococci earlier in life and suffer the greatest burden of pneumococcal disease (3). High density colonization is associated with an increased likelihood of pneumococcal disease in children, and transmission in experimental models (4–8).

Pneumococcal Conjugate Vaccines (PCVs) in widespread use for children target between 10-13 pneumococcal serotypes by inducing an antibody response to the capsular polysaccharide. PCVs have been an outstanding public health success, providing direct protection against both carriage and disease, as well as indirect (herd) protection in some settings. Vaccinated individuals are less likely to transmit pneumococci to unvaccinated contacts, reducing acquisition (9).

Despite the success of PCVs, drawbacks include their high cost, protection against a limited number of serotypes, and their potential to drive serotype replacement (an increase in non-vaccine serotypes in carriage and disease in vaccinated populations). In addition, immune responses to pneumococcal vaccines are reduced in children who are colonized by pneumococci at the time of vaccination (10–13). Vaccine-induced IgG levels are reduced in carriers compared with non-carriers (10, 13). This hypo-responsiveness is serotype-dependent, with reduced serotype-specific IgG levels occurring in response to PCV when the homologous serotype was carried at the time of vaccination (11, 12). Children in high burden settings are more likely to be colonized at the time of vaccination, which occurs within the first year of life (3). For example, longitudinal sampling of a South African cohort of infants identified the median age of first pneumococcal acquisition was 63 days old (14). Similar studies in Thailand (15) and Indonesia (16) found the median age of acquisition was 45.5 and 129 days old, respectively. As such, the immune responses to PCV might not be optimal in these groups.

To combat some of the limitations associated with currently licensed PCVs, alternative pneumococcal vaccines are under investigation. These include expanded valency PCVs, protein-based vaccines, and whole cell vaccines (17). Whole cell vaccines comprise an inactivated pneumococcal strain. Research into whole cell vaccines predates the use of capsular vaccines, with the some of the earliest pneumococcal whole vaccine trials conducted by Sir Almroth Wright in the early 1900s, administering a heat-killed preparation to gold miners in South Africa (18). Following the discovery of serotype specificity, determined by capsule type, attention then turned to polysaccharide-based vaccines. Only in recent years has pneumococcal whole cell vaccines once again garnered attention.

In the early 2000s, investigations began into an ethanol killed whole cell vaccine, which was able to prevent colonization and disease in animal models (19). This vaccine differed from the vaccine used in the 1900s by both the method of inactivation and the strain used. The use of a non-encapsulated strain provided a simple strategy to allow for greater exposure of pneumococcal surface antigens to the host. The pneumococcal strain used, RX1AL-, is a spontaneous capsule-negative mutant derived from well-known laboratory strain D39 with a deletion of the *lytA* gene (20) to enable high-density growth without bacterial autolysis. After demonstrating powerful serotype-independent protection against both pneumococcal colonization and invasive disease in pre-clinical models, the whole cell vaccine preparation was modified for use in humans. Subsequent studies employed a modified RX1AL-strain (termed RM200) in which the pneumolysin gene had been altered to express a non-haemolytic derivative of Ply (21). In its current formulation, this whole cell vaccine (hereby referred to as WCV) consists of beta-propiolactone inactivated RM200 adsorbed to aluminium hydroxide adjuvant. WCV was shown to be safe and well tolerated in rabbit toxicity studies and accelerated the clearance of pneumococcal colonization in adult mice when given prior to pneumococcal challenge (22). This accelerated clearance required CD4+ T cells and IL-17A signalling, but antibody-independent (22–24).

Using a preventative model, we previously found that while a single dose of WCV given to infant mice resulted in lower pneumococcal density in the nasopharynx and middle ear compared with unvaccinated mice, it did not prevent middle ear infection. The reduction of pneumococcal density in the middle ear required both antibodies and CD4+ T cells in this model (25). This WCV has also been tested in a phase 1 trial, where it was found to be safe and immunogenic in healthy adults (26). More recently, research into a γ-irradiated whole cell vaccine has commenced, which protected mice from pneumococcal pneumonia and sepsis (27–29) and also elicited opsonophagocytic killing of encapsulated pneumococci (30).

Given that a large proportion of infants in high disease burden settings carry pneumococci at the time of vaccination, our aim in this study was to determine how prior pneumococcal colonization affects WCV-induced immunity, and the impact of this ‘therapeutic immunization’ on nasopharyngeal colonization, shedding, and middle ear infection in a murine model.

## Results

To establish a long-term pneumococcal colonization model we used infant mice to reflect the carriage dynamics observed in young children. Intranasal infection of five-day old mice with 2 x 10^3^ CFU of pneumococcal strain EF3030 (serotype 19F) led to rapid *in vivo* replication resulting in localized carriage in the nasopharynx, which was maintained at a consistent density of 10^5^-10^6^ CFU for up to 45 days post-infection. After 45 days, colonization gradually reduced until cleared 65 days post-infection (Figure 1A).

**Figure 1:**
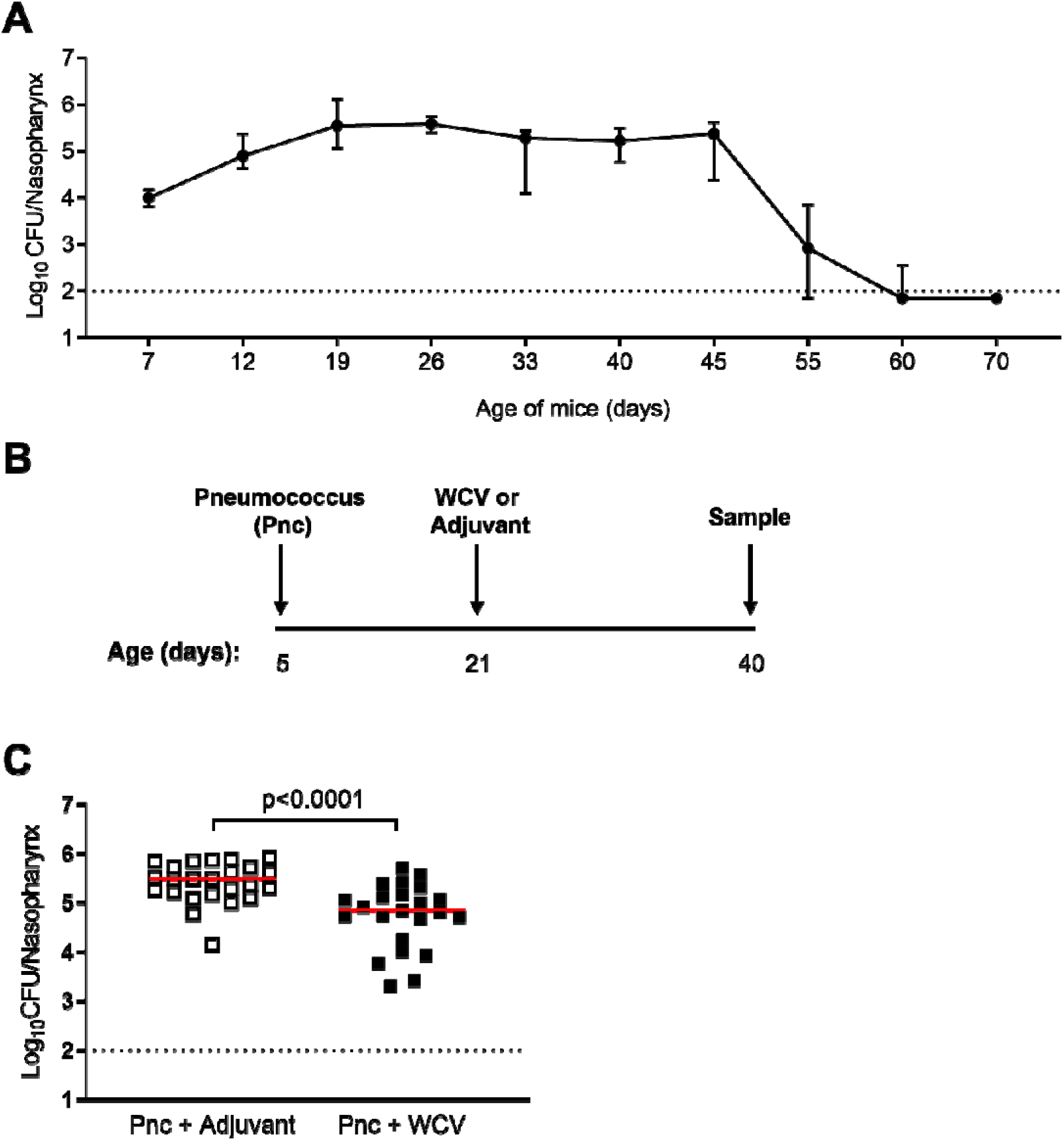
Effect of therapeutic pneumococcal whole cell vaccine (WCV) on pneumococcal colonization density. A) Pneumococcal colonization density in nasopharyngeal tissues following intranasal infection of five-day-old mice with 2 x 10^3^ CFU of strain EF3030 (serotype 19F). Data presented as the median ± interquartile range with ∼10 mice per timepoint. B) Schematic of therapeutic WCV model. Five-day-old mice receive 2 x 10^3^ CFU of strain EF3030 via intranasal administration. At 21 days old, WCV or adjuvant was given subcutaneously. Mice were euthanized at 40 days old for tissue collection. C) Pneumococcal nasopharyngeal density in mice given therapeutic adjuvant (open squares) or WCV (closed squares). Solid red lines denote the median and the black dotted line represents the assay limit of detection. Statistics: Mann Whitney U test, with p-values < 0.05 shown.

To determine the effect of therapeutic immunization of WCV on pneumococcal carriage, we infected five-day old mice with EF3030 to establish pneumococcal colonization and then vaccinated mice 16 days later (at 21 days old), at which point mice were experiencing high density (∼10^5^) carriage (Figure 1B). Mice were vaccinated with a single subcutaneous dose of WCV (beta-propiolactone inactivated pneumococci RM200 with aluminium hydroxide adjuvant, “therapeutic WCV”) or adjuvant only as a control (“therapeutic adjuvant”). Pneumococcal density in the nasopharynx was measured when mice were 40 days old (35 days post-pneumococcal infection). Therapeutic administration of WCV resulted in a 0.63 log_10_ reduction in nasopharyngeal density compared with mice given therapeutic adjuvant alone (p < 0.0001, Mann Whitney U test, Figure 1C).

We next investigated the immunological mechanisms mediating this therapeutic effect. In adult mice, IL-17A responses mediate WCV-induced protection against pneumococcal colonization, and pre-infection concentrations of *ex vivo* elicited IL-17A in whole blood are inversely correlated with post-infection pneumococcal colonisation density (24). To investigate if IL-17A responses were involved in the observed therapeutic WCV-induced reduction in colonization, we measured IL-17A concentrations in nasopharyngeal tissues, sera, and in splenocytes stimulated *ex vivo* with WCV antigen. There were no differences in IL-17A responses in nasopharyngeal tissue and sera (Figure 2A) or in splenocytes stimulated with WCV antigen (Figure 2B) between mice given therapeutic WCV or therapeutic adjuvant.

**Figure 2:**
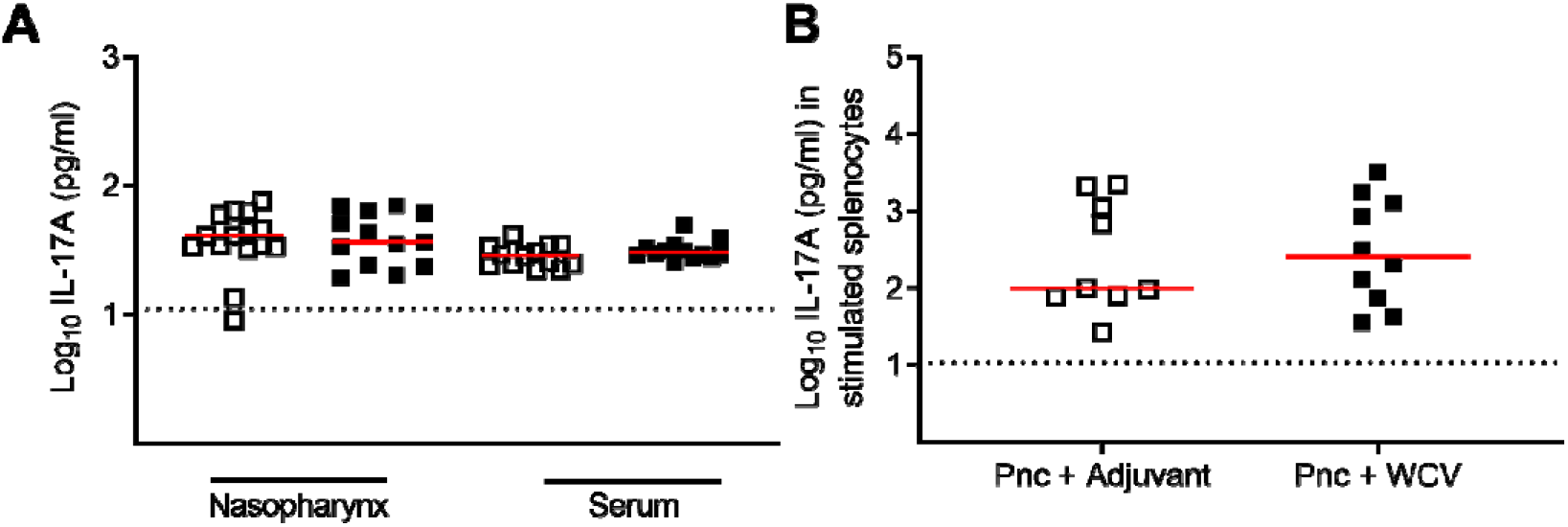
IL-17A concentrations in mice given therapeutic adjuvant or WCV. A) IL-17A concentrations in nasopharyngeal tissue homogenates and sera from mice given therapeutic adjuvant (open squares) or WCV (closed squares) as described in Figure 1B. B) IL-17A concentrations in splenocyte suspensions following *in vitro* stimulation with WCV antigen. Solid red lines denote the median and the black dotted line represents the assay limit of detection. Statistics: Mann Whitney U test, with p-values < 0.05 shown.

We next investigated the role of antibodies in exerting the therapeutic effect on pneumococcal carriage. Therapeutic administration of WCV led to a 1.6 log_10_ increase in WCV-specific IgG levels (defined as any non-capsular antibodies induced in response to WCV antigens) in plasma compared with the adjuvant control (p < 0.0001, Mann Whitney U test, Figure 3A). To confirm that antibodies were responsible for the reduction in colonization density, we colonized and vaccinated antibody-deficient C57BL/6.μMT^-/-^ mice using the same timelines as Figure 1B. In contrast with C57BL/6 mice (Figure 1C), no difference in pneumococcal density was observed in μMT^-/-^ mice were given therapeutic WCV compared to adjuvant (p = 0.18, Mann Whitney U test, Figure 3B).

**Figure 3:**
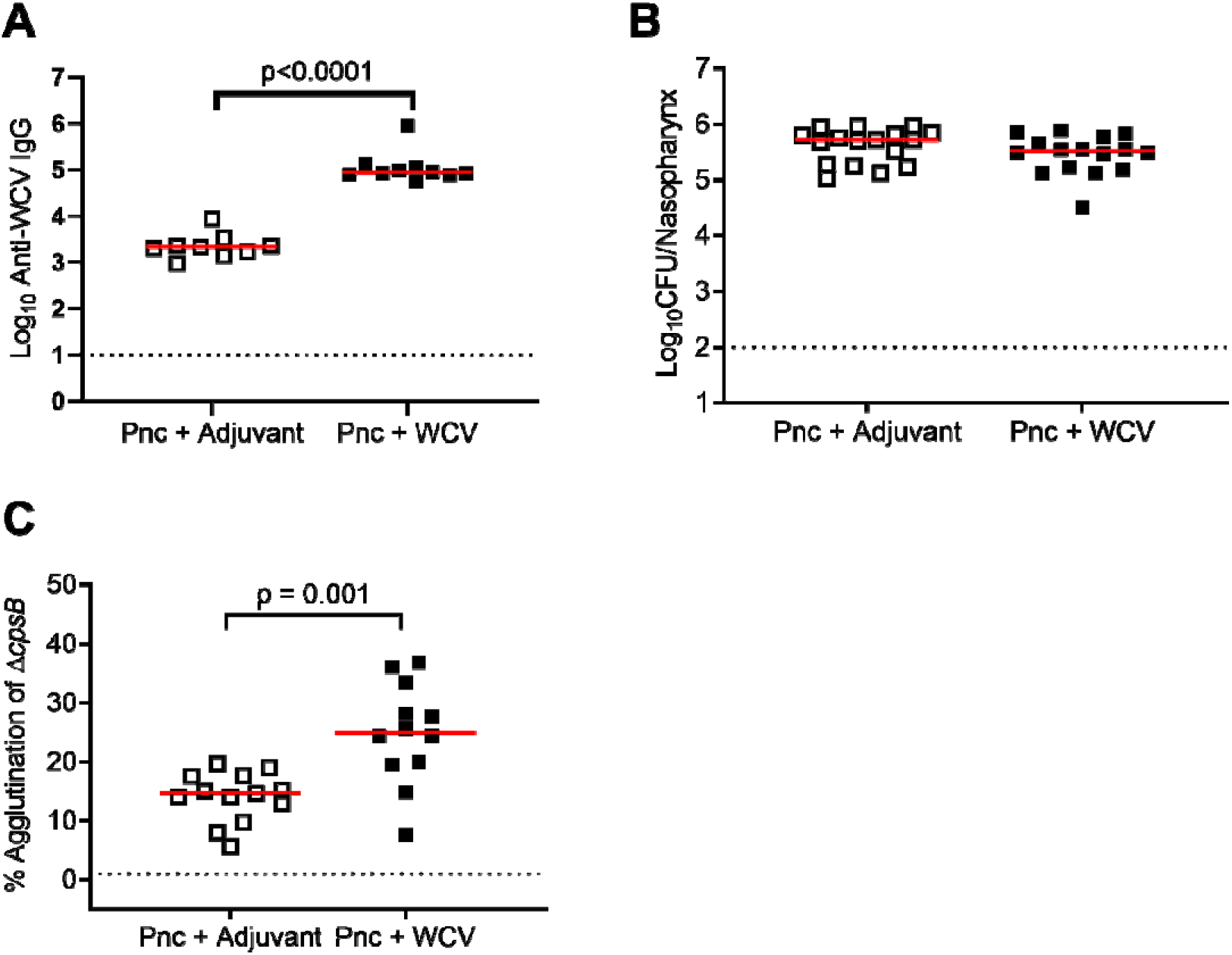
Effect of therapeutic WCV on vaccine-specific IgG responses. A) Anti-WCV IgG in plasma of mice given therapeutic adjuvant (open squares) or WCV (closed squares) as described in Figure 1B. B) Pneumococcal nasopharyngeal density in antibody-deficient (µMT^-/-^) mice colonized by pneumococci and vaccinated as described in Figure 1B. C) Agglutination percentage of pneumococcal capsule deficient mutant Δ*cpsB* upon incubation with sera from mice given therapeutic adjuvant or WCV. Solid red lines denote the median and the black dotted line represents the assay limit of detection. Statistics: IgG and density data were analyzed using the Mann Whitney U test, with p-values < 0.05 shown; agglutination data were analyzed using Welch’s t-test

Given the agglutinating activity of antibodies contributes to prevention as well as reduction in pneumococcal colonization density (31–33), we considered whether the reduction in pneumococcal density following therapeutic WCV was attributable to the agglutinating ability of anti-WCV antibodies. Sera from colonized mice that received therapeutic WCV or adjuvant were incubated with Δ*cpsB,* a capsule deficient mutant of EF3030 and analyzed by flow cytometry. When Δ*cpsB* was incubated with sera from mice given therapeutic WCV, there was a 11% increase in agglutination compared with the adjuvant control (Figure 3C, p = 0.001, Welch’s t-test). Taken together, our data suggest that therapeutic administration of WCV induces antibody responses with agglutinating abilities that mediate a reduction in colonization density.

To identify some of the specific pneumococcal antigens that were targeted by anti-WCV antibodies, we analysed plasma from therapeutic WCV and adjuvant mice for IgG directed against 27 pneumococcal proteins using a direct binding multiplex assay (34). We found that therapeutic WCV significantly enhanced IgG levels to three (CbpA, PiaA and PspA [family 1]) of 27 the proteins tested, compared with therapeutic adjuvant (Figure 4A and Supplementary Table 1).

**Figure 4:**
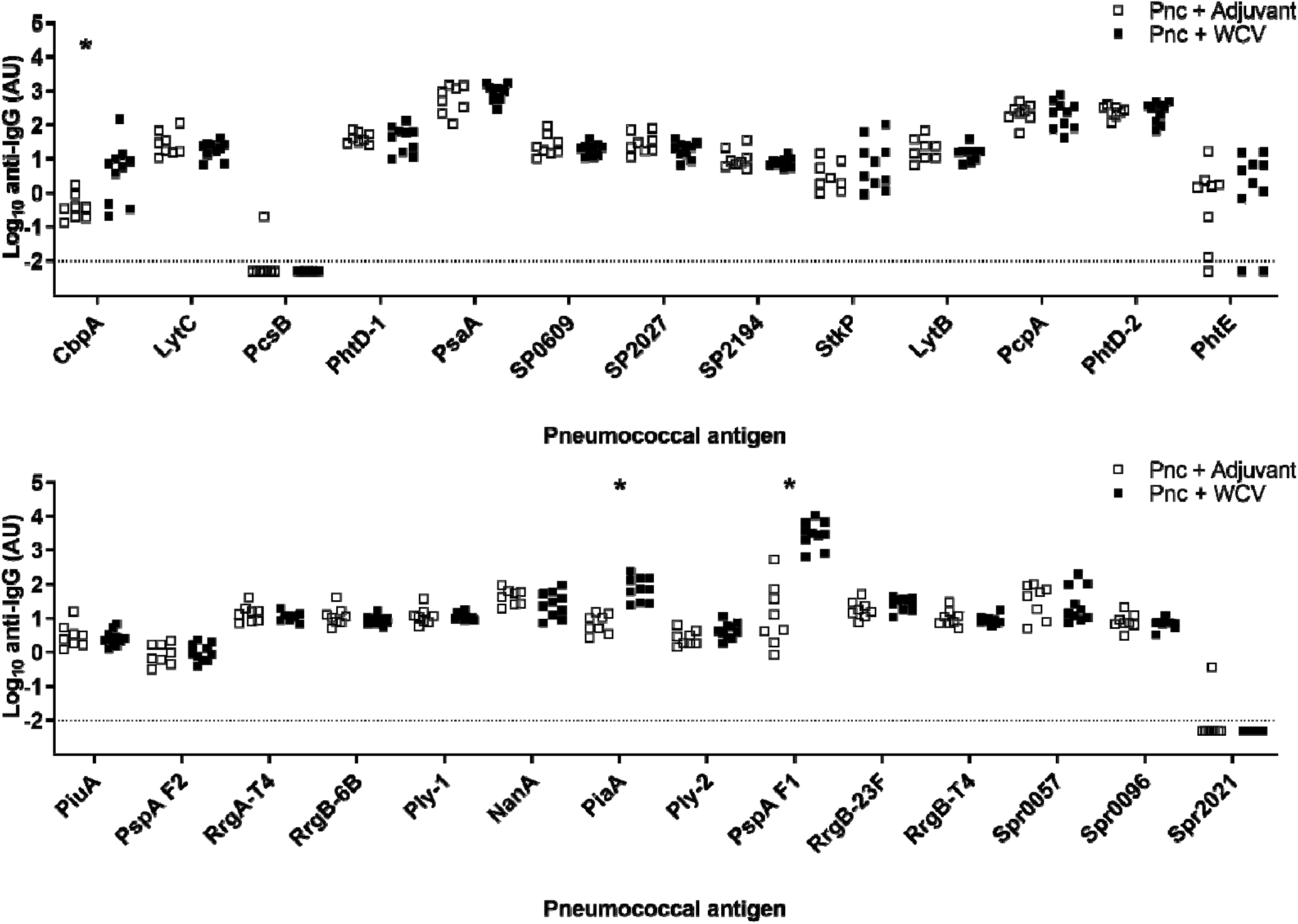
Immune response to non-capsular antigens elicited by therapeutic WCV or adjuvant administration. IgG responses to pneumococcal surface antigens in plasma following therapeutic adjuvant (open squares) or WCV (closed squares) administration. Dotted line represents the assay limit of detection. RrgA and RrgB are subunits of pneumococcal pilus-1, which is not present in either the WCV or pneumococcal EF3030 strain used in this study. Ply-1 and Ply-2 are different variants of pneumolysin. Statistics: IgG data were analyzed by using the Mann Whitney U test, with p values < 0.05 denoted by *

Because pneumococcal nasopharyngeal density is associated with disease, we hypothesized that therapeutic WCV would be able to prevent infection of the middle ear of infant mice. To test this, we induced pneumococcal middle ear infection by co-infection with influenza A virus (IAV) to facilitate bacterial dissemination to this site. Mice were colonized in the nasopharynx with pneumococci at 5 days old, given a single dose of WCV or adjuvant at 21 days of age, co-infected with IAV at 28 days of age, after which pneumococcal density was measured in nasopharyngeal and middle ear tissues at 40 days of age (Figure 5A). Mice receiving therapeutic WCV exhibited a 0.62 log_10_ (p = 0.0007, Mann Whitney U test) and 1.45 log_10_ (p = 0.0019, Mann Whitney U test) reduction in pneumococcal density in both the nasopharynx and middle ear, respectively, compared with mice that received therapeutic adjuvant (Figure 5B). Additionally, mice given therapeutic WCV were less likely to have bilateral infection, with 4/15 (26.7%) therapeutic WCV mice having detectable pneumococci in both middle ears compared with 10/14 (71.4%) of mice given therapeutic adjuvant (p = 0.027, Fisher’s exact test). By contrast, there was no difference between experimental groups for mice exhibiting no or unilateral middle ear infection (Table 1).

**Figure 5:**
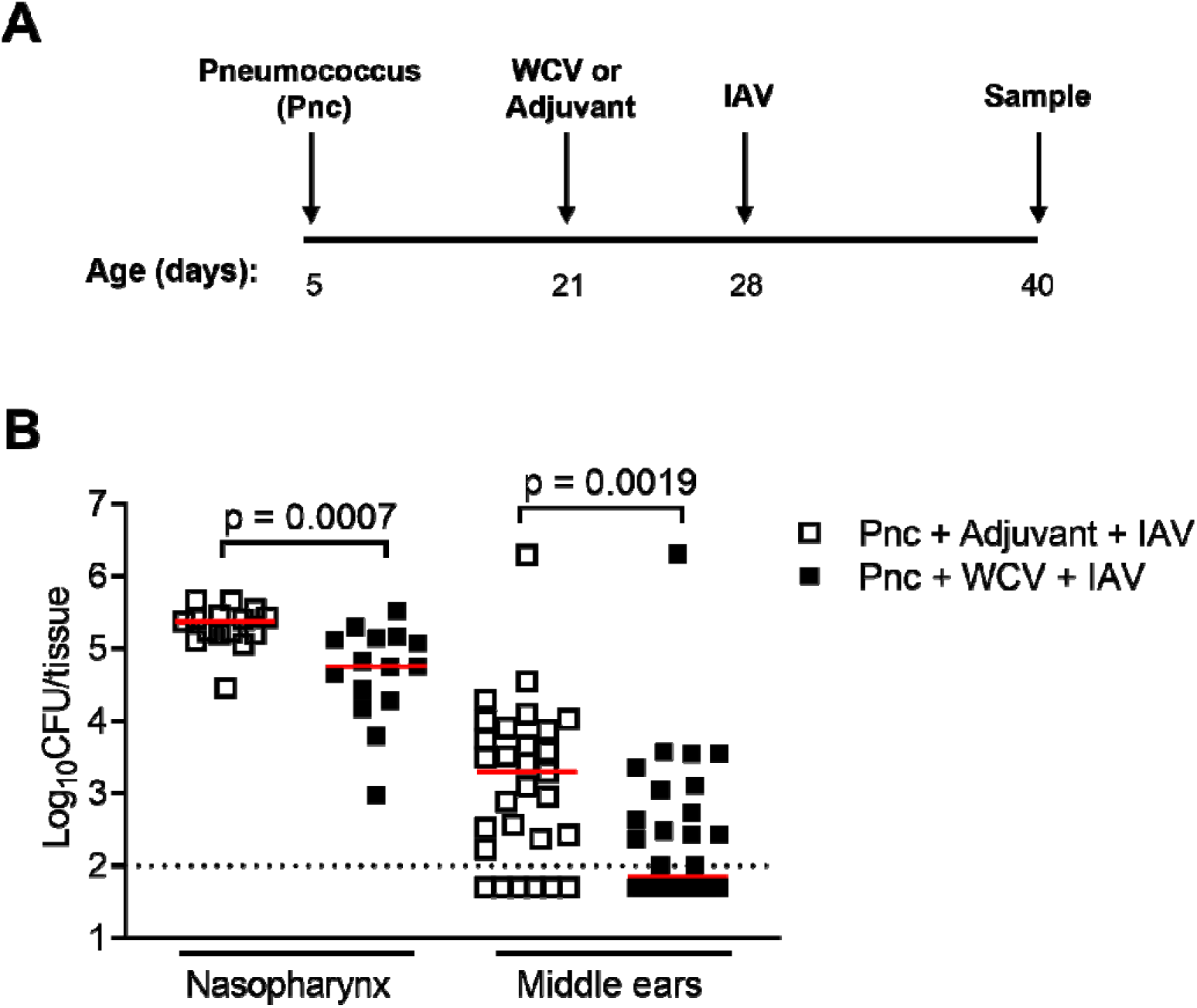
Effect of therapeutic WCV on influenza A virus (IAV)-induced pneumococcal infection of the middle ear. A) Schematic representation of pneumococcal middle ear infection model. Five-day-old mice received 2 x 10^3^ CFU of pneumococcal strain EF3030 via intranasal administration, followed by subcutaneous immunization with WCV or adjuvant at 21 days of age. At 28 days old, mice were infected with 300 PFU IAV (strain A/Udorn/72) to facilitate pneumococcal dissemination to the middle ear. Mice were euthanized at 40 days old for tissue collection. B) Pneumococcal density in the nasopharynx and middle ears of mice given therapeutic adjuvant (open squares) or WCV (solid squares). Each mouse has two data points for middle ear samples (one for each middle ear). Solid red lines denote the median and the black dotted line represents the assay limit of detection. Statistics: Mann Whitney U test, with p-values < 0.05 shown.

**Table 1.** Effect of therapeutic WCV on pneumococcal OM.

| OM type | Pnc + Adjuvant,<br>Number of mice (%) | Pnc + WCV, Number<br>of mice (%) | p value<sup>a</sup> |
| --- | --- | --- | --- |
| No OM | 2/14 (14.3) | 4/15 (26.7) | 1 |
| Unilateral | 2/14 (14.3) | 7/15 (46.7) | 0.11 |
| Bilateral | 10/14 (71.4) | 4/15 (26.7) | **0.027** |

<sup>a</sup>Comparison of Pnc + adjuvant vs Pnc + WCV, Fisher's exact test. Significant findings ( $p < 0.05$ ) are shown in bold.

Given the importance of pneumococcal density in shedding and transmission, we hypothesized that therapeutic WCV administration might also reduce pneumococcal shedding, which we measured by gently tapping the nose of mice onto selective media for pneumococcal culture and enumeration, expressed as CFU/10 taps. Preliminary data revealed that therapeutic WCV reduced pneumococcal shedding in mice five days after immunization, but pneumococcal density rebounded by ten days after immunization (Figure 6A). To investigate whether the effect of therapeutic WCV on shedding could be prolonged, we tested a two-dose vaccine schedule (Figure 6B). Mice were colonized with pneumococci at five days of age and vaccinated with either WCV or the adjuvant control at 14 and 28 days old, followed by euthanasia and sample collection at 40 days old. This therapeutic WCV immunization schedule reduced pneumococcal density by 0.94 log_10_ compared with therapeutic adjuvant (p = 0.014, Mann Whitney U test, Figure 6C).

**Figure 6:**
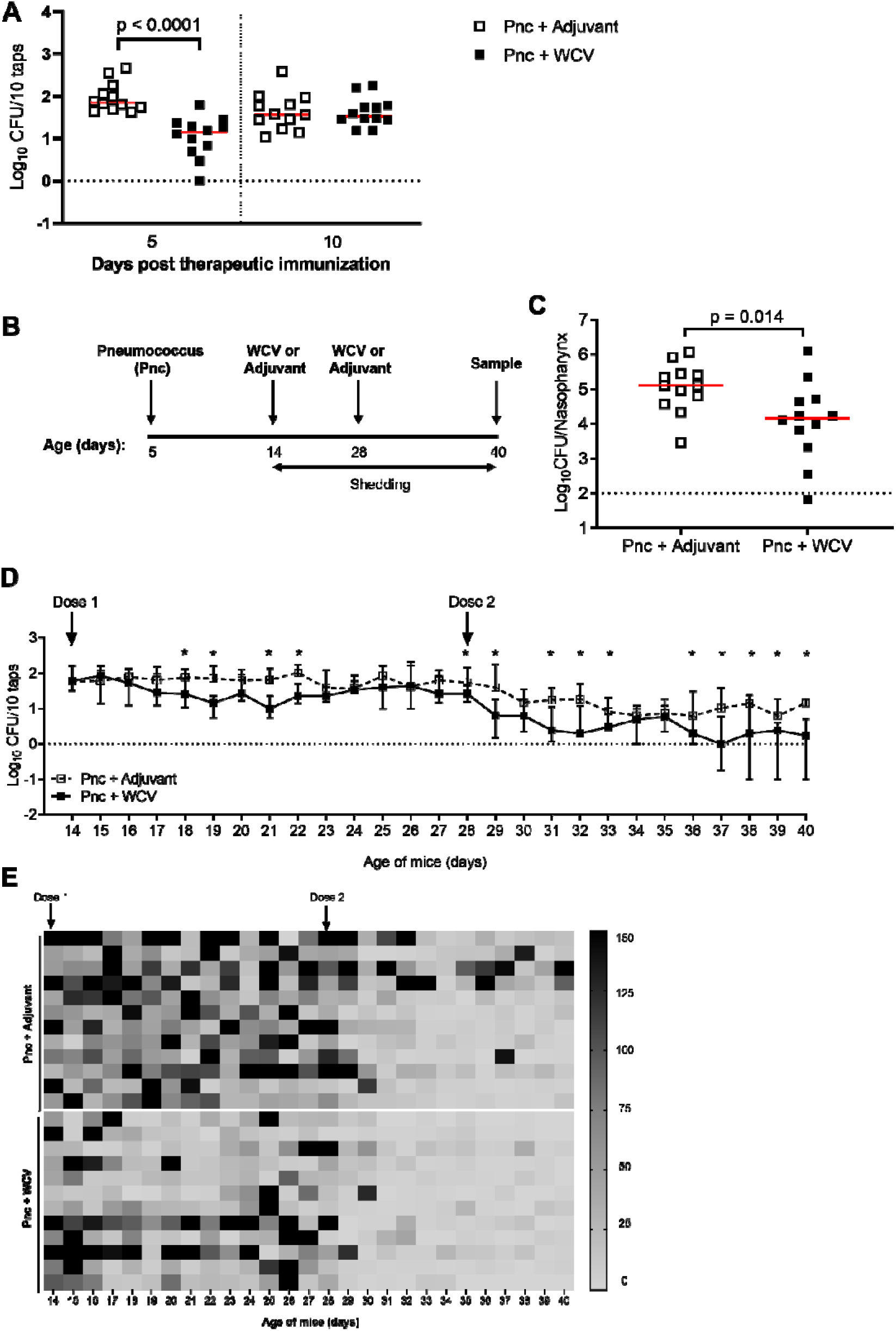
Effect of therapeutic WCV on pneumococcal shedding. A) Pneumococcal shedding in mice receiving one therapeutic dose of adjuvant (open squares) or WCV (closed squares) measured at 5- and 10-days post-immunization. B) Schematic representation of two dose-immunization model for therapeutic vaccination. Five-day-old mice receive 2 x 10^3^ CFU of pneumococcal strain EF3030 via intranasal administration, followed by subcutaneous immunization with WCV or adjuvant at 14 and 28 days of age. Pneumococcal shedding was measured daily from 14-40 days of age. Mice were euthanized at 40 days old for tissue collection to examine pneumococcal density. C) Pneumococcal nasopharyngeal density in 40-day old mice given therapeutic adjuvant (open squares) or WCV (closed squares) in the two-dose schedule described in Figure 6B. Solid red lines denote the median and the black dotted line represents the assay limit of detection. D) Pneumococcal shedding measured in mice given therapeutic adjuvant or WCV in a two-dose schedule (14 and 28 days old) with 12 mice in each group. Data are presented as the median with the interquartile range. E) Heat map of the number of pneumococci shed by individual mice over time, where each row represents an individual mouse and high shedding events (defined as ≥ 150 colonies/10 taps) are shown in red. Statistics: Mann Whitney U test, with p values < 0.05 shown or denoted by *.

After establishing the two-dose schedule, we measured pneumococcal shedding daily as described above (Figure 6D). Following the first dose of WCV, a reduction in shedding was observed from 4-8 days post-WCV compared with adjuvant, which rebounded between 9-13 days. Following the second dose of WCV, a more pronounced reduction in pneumococcal shedding occurred with significant reductions observed at days 0-5 and 8-12 after the second WCV dose compared with adjuvant.

Previous studies have demonstrated that mice shedding > 300 CFU/20 taps are significantly more likely to transmit pneumococci to co-housed infant mice (35). We therefore analyzed our data stratified by the number of pneumococci shed in individual mice over time and defined a “high shedding event” as ≥ 150 colonies/10 taps (denoted as black in Figure 6E). We observed fewer high shedding events in mice given WCV compared with mice given adjuvant (Figure 6E: 29/324 [9%] vs. 47/324 [14.5%], p = 0.037, Fisher’s exact test). Furthermore, 0/144 high-shedding events were observed following the second dose of WCV, whereas as 9/144 (6%) high shedding events occurred following a second dose of adjuvant (p = 0.003, Fisher’s exact test) (Figure 6E).

## Discussion

Although PCVs prevent pneumococcal transmission and disease, their limitations around serotype replacement and hypo-responsiveness in children carrying pneumococci have sparked interest in alternative strategies. In this study, we examined a serotype-independent whole cell vaccine (WCV) and showed that when given therapeutically to mice, WCV reduced pneumococcal carriage density, shedding and middle ear infection.

When we explored the mechanisms by which therapeutic WCV administration reduced nasopharyngeal carriage density, it was evident that there was no association with IL-17A concentrations (Figure 2). This was unexpected as IL-17A is known to control pneumococcal carriage density and duration (24, 36) and preventative administration of the same WCV promoted IL-17A production and pneumococcal clearance from the nasopharynx (22). The latter study administered 2-3 doses of WCV to adult mice prior to challenge whereas our study administered 1-2 doses of WCV to infant mice after challenge. Additionally, it is also possible that the timepoint we measured IL-17A concentrations was too late in the infection to observe any association. Together, these differences might explain the observed discrepancies in IL-17A responses, particularly the use of infant mice, as data from mice and humans indicate that Th17 and IL-17A responses increase with age (37, 38). Additional experiments using mice deficient in IL-17A or its receptor might confirm the role of this cytokine in our model.

In this study, the reduction in pneumococcal carriage density mediated by therapeutic administration of WCV was antibody-dependent, demonstrated by no reduction in density when therapeutic WCV was given to antibody-deficient mice (Figure 3B). Antibodies are involved in the prevention of pneumococcal carriage as well as mediating clearance from the nasopharynx through agglutination mechanisms (31, 39, 40). In our study, it is not clear whether the enhanced agglutination following therapeutic WCV is due to capsular or non-capsular antibodies as the agglutination assays were conducted with a mutant of EF3030 (Δ*cpsB*) that was deficient (but not incapable of) producing capsule (41). However, the WCV strain used in this study is non-encapsulated, making it unlikely enhanced agglutination in therapeutic WCV mice would be caused by capsular antibodies. Nonetheless, it is evident that therapeutic administration of WCV enhances the production of agglutinating antibodies that reduce pneumococcal carriage density.

Therapeutic WCV induced a robust humoral response against three non-capsular antigens on the cell surface of pneumococci; CbpA, PspA (family 1) and PiaA. A phase 1 trial in adults with the same WCV identified increased IgG levels in some pneumococcal antigens including PspA (26), which is consistent with our study. In contrast, the trial identified elevated IgG levels to Ply. The reason for the differences in WCV-induced IgG levels to Ply is unclear.

Both PspA and CbpA are important in nasopharyngeal colonization and thus have been previously identified as vaccine candidates (42), but are highly diverse among pneumococci. PiaA has also been previously explored as a vaccine candidate and is highly conserved (43). A challenge in assessing immunogenicity of whole cell vaccines is that responses to individual antigens can be heterogeneous. In our study, antibody responses in WCV-immunized mice were consistently high for PspA and PiaA, but varied for CbpA. Additionally, only 27 pneumococcal antigens were examined in our study. It is therefore unclear which antigens (including those not examined in this study) are important contributors to the reduction in pneumococcal carriage density observed in our model.

One of the major benefits of PCVs is their ability to reduce carriage of vaccine serotypes, which reduces transmission, resulting in herd protection (9, 44). Shedding from a colonized individual is an important step in murine transmission models, with previous studies identifying a shedding ‘threshold’ required for transmission to occur (35). We found that WCV decreased shedding to below this threshold (Figure 6D and 6E), suggesting it may reduce pneumococcal transmission. The basis of the reduced shedding mediated by WCV is unclear, but likely due to the reduction in pneumococcal density in the nasopharynx (41, 45).

Pneumococcal carriage density is also important in disease. Higher pneumococcal carriage density is observed in individuals with pneumococcal disease compared with healthy individuals (6–8). We found that therapeutic WCV administration reduced pneumococcal infection of the middle ear in a co-infection model (Figure 5B). Previously, we showed that WCV given in a preventative manner (6 days before pneumococcal challenge) reduced pneumococcal density in the middle ear but not in the nasopharynx (25). Interestingly, prophylactic administration of WCV did not reduce bilateral middle ear infection (25) whereas therapeutic WCV administration did (Table 1). This difference could be explained by differences in the immune response to WCV either due to mode of administration (therapeutic vs preventative) or the age at which mice were vaccinated.

Our study is subject to several limitations, such as the use of a single pneumococcal strain. Additionally, we tested the potential for therapeutic vaccination to reduce non-invasive disease. In future studies, it would be of interest to test if therapeutic WCV administration can reduce invasive pneumococcal disease. Lastly, as with all pre-clinical models, it is unclear how our findings translate to humans. However, our study provides valuable data to suggest there is merit in investigating the effects of WCV when administered therapeutically in humans.

In this study, we showed that giving WCV to mice already colonized by pneumococci can reduce pneumococcal carriage density, shedding and middle ear infection. Our findings could help inform the development and evaluation of new pneumococcal vaccines that includes consideration of the effects of vaccination when given to an infant already colonized by pneumococci. Our data suggest the use of vaccines in children with similar properties may be beneficial in these settings in reducing carriage density, transmission, and disease.

## Acknowledgements

We thank staff at the Biological Resources Facility, the University of Melbourne at the Peter Doherty Institute for Infection and Immunity, and the Disease Model Unit, the Murdoch Children’s Research Institute, for their expert assistance in animal husbandry. We thank Professors Lorena Brown and Frank Carbone (Department of Microbiology and Immunology, The University of Melbourne at the Peter Doherty Institute for Infection and Immunity) for IAV A/Udorn/72/307 and mouse μMT^-/-^, strains respectively. We also thank Polly Burbidge for advice and expertise regarding the MSD platform.

This study was supported by a Robert Austrian Research Award in Pneumococcal Vaccinology awarded to JM by the International Symposium on Pneumococci and Pneumococcal Diseases (ISPPD, funded by Pfizer) as well as a National Health and Medical Research Council (NHMRC) Ideas Grant (GNT1182442) and Infection and Immunity theme investment funds from the Murdoch Children’s Research Institute. CS was a recipient of a NHMRC Career Development Fellowship (GNT1087957). This work was also supported by the Victorian Government’s Operational Infrastructure Support Program.

## Author contributions

Conceptualization: JM, EMD, EKM, PL, RRB, RM, CS, OW

Methodology: JM, SM, EMD, PB, DG, PL, RRB, RM, CS, OW

Formal analysis: JM, SM, CS, OW

Investigation: JM, SM, VB, CLP, NP, SK, PD, PB, DG

Resources: EKM, RM, DG, CS, OW, PL

Funding acquisition: JM, SM, EKM, CS, OW

Writing of the original manuscript draft: JM, SM, CS, OW

Review and editing of the manuscript for submission: JM, SM, EMD, VB, CLP, NP, SK, PD, PB, DG, EKM, PL, RRB, RM, CS, OW

## Methods

### Bacterial and viral strains

EF3030 is a serotype 19F pneumococcal strain originally isolated from a patient with otitis media (46). Bacteria were grown in 3% (w/v) Todd Hewitt Broth (Oxoid) supplemented with 0.5% (w/v) yeast extract (Becton Dickinson) or on Horse Blood Agar (HBA, Oxoid) at 37°C in 5% CO_2_. Pneumococcal and Influenza A virus (IAV) strain A/Udorn/72/307 stocks for mouse experiments were prepared as previously described (47).

### Vaccine preparations

Whole cell antigen (WCA) comprised lyophilized pneumococcal strain RM200 killed by beta-propiolactone (BPL)-treatment. RM200 is a derivative of the unencapsulated pneumococcal strain RX1E, with a deleted autolysin gene and a non-toxic pneumolysoid (48). To obtain working stocks, WCA was resuspended in sterile dH_2_O to 10.8 mg/ml and stored at -20°C. To prepare WCV (49), WCA was incubated with aluminium hydroxide (Alhydrogel, Brenntag) overnight at 4°C with rotation. Each 100 μl WCV dose contained 100 μg of WCA and 2.5 μg/μl Alhydrogel in sterile saline. In all immunization experiments, the effects of WCV were compared to a group of mice receiving the same dose of Alhydrogel in 100 μl of saline (referred to as adjuvant throughout).

### Study approval

All animal experiments in this study were approved by the Animal Ethics Committee of the University of Melbourne (AEC1413144) or the Animal Ethics Committee at the Murdoch Children’s Research Institute (A847, A851), and were conducted in strict accordance with the Prevention of Cruelty to Animals Act (1986) and the Australian National Health and Medical Research Council Code of Practice for the Care and Use of Animals for Scientific Purposes (2013).

### Mice

C57BL/6 and C57BL/6.μMT^-/-^ (50) mice were bred and housed at the Department of Microbiology and Immunology at The Peter Doherty Institute for Infection and Immunity, The University of Melbourne, Australia or the Murdoch Children’s Research Institute. Litters of mice were housed in ventilated cages with unlimited access to food and water. Pups were weaned at 21 days of age with males and females placed into separate cages with a maximum of five mice per cage.

### Murine models

To establish colonization, litters of five-day-old C67BL/6 were infected intranasally without anaesthesia with 2 x 10^3^ CFU of pneumococci in 3 μl of PBS.

Mice were immunised with WCV or adjuvant via subcutaneous injection to the back of the neck at either 21 days old (one dose schedule) or at 14 and 28 days old (two dose schedule).

To induce pneumococcal infection of the middle ear, mice intranasally infected with pneumococci as described above, were subsequently infected with 300 PFU of IAV given intranasally in 3 μl of PBS at 28 days of age (25).

Initially, mice were euthanized by CO_2_ inhalation. However, following updates in animal handling practices, mice were subsequently anesthetized using isoflurane and then euthanized by cervical dislocation at the time points specified. Method of euthanasia did not result in any changes in the model. Nasopharyngeal and middle ear tissue was collected for enumeration of pneumococci by viable count on HBA plates supplemented with 5 µg/ml gentamicin. Terminal blood samples and spleen were harvested and analysed for immune responses.

Measurement of pneumococcal shedding was conducted as described previously (35, 51) with the following modifications. The nose of each mouse was gently tapped 10 times across two HBA plates supplemented with 5 µg/ml gentamicin (selective for pneumococci) and spread across the plates using a 10 µl disposable loop dipped in PBS.

### Immune responses

To perform *ex vivo* stimulation, whole blood and splenocyte suspensions were stimulated with whole cell antigen (WCA) or PBS (as a control). Whole blood was diluted 1:10 in DMEM/F12 and added to duplicate wells of a 24-well tray. WCA was added to one well at a final concentration of 10^7^ CFU/ml and the same volume of sterile PBS was added to the remaining well as a negative control. For splenocyte stimulation, single cell suspensions of splenocytes were seeded into duplicate wells of a 24 well tray at a concentration of 10^7^ cells/ml. WCA was added to one well at a final concentration of 10^6^ CFU/ml, and the same volume of PBS added to the remaining well as a negative control. Stimulated whole blood and splenocytes were incubated for 72 h at 37°C and stored at -80°C until tested. IL-17A was measured in nasopharyngeal homogenates, sera and culture supernatants from stimulated splenocytes using the Quantikine Mouse IL-17 ELISA kit (R&D Systems Inc.) according to the manufacturer’s instructions. WCA-specific IgG was detected in plasma or in whole blood following *ex vivo* stimulation using the method described previously (22) with some modifications (25).

IgG responses to pneumococcal surface antigens in plasma following therapeutic adjuvant or WCV administration were measured using the MSD platform as described previously (34, 52).

### Agglutination assay

Agglutination assays were performed using a method described previously (33, 53). Pneumococcal infectious stocks (as described above) of *ΔcpsB* capsule-deficient strain were thawed, washed (11,300 x g for 5 min) and diluted in PBS. To each well, 10^5^ CFU of pneumococci and 25 μl of serum were added and incubated for 1.5 h at 37°C and 5% CO_2_. Bacteria were then fixed with 1% (v/v) paraformaldehyde (Sigma Aldrich) diluted 1:2 with PBS and analysed by flow cytometry (LSR Fortessa X-20, BD Biosciences) using FacsDiva Software. Agglutination of pneumococci was visualised as a shift in forward scatter using dot plots which corresponds to increasing particle size. Agglutination in response to different sera was quantified by calculating the percentage of bacteria with an altered forward scatter compared to bacteria alone. Background produced by sera alone was subtracted from individual samples.

### Statistics

Data obtained from experimental groups that were not normally distributed were compared using the Mann Whitney U Test. Data that were normally distributed were analyzed by Welch’s t-test. Fisher’s exact test was used to compare proportion differences. The statistical test used for each analysis is indicated in the accompanying text and figure legends. P values less than 0.05 were considered statistically significant.

