## Supplementaty Table1 for "Therapeutic immunization with a whole cell vaccine reduces pneumococcal nasopharyngeal density, shedding, and middle ear infection in mice"

### Supplemental information titles and legends

**Table S1.** Therapeutic effects of WCV on antibody responses to pneumococcal proteins.

|  | **Median IgG AU (95% CI)** | | | | |
| --- | --- | --- | --- | --- | --- |
| **Antigen** | **19F + Adjuvant** | | **19F + WCV** | | **p value**^a^ |
| CbpA | 0.4 | (0.1, 1.7) | 6.6 | (0.3, 13.2) | **0.012** |
| LytC | 24.2 | (10.9, 113.7) | 22.6 | (7.4, 28.7) | > 0.05 |
| PcsB | 0.0 | (0.0, 0.2) | 0.0 | (0, 0) | > 0.05 |
| PhtD-1 | 38.4 | (26.0, 73.6) | 53.5 | (11.7, 86.4) | > 0.05 |
| PsaA | 725.9 | (110.8, 1512) | 1055 | (545.3, 1640) | > 0.05 |
| SP0609 | 20.3 | (10.3, 95.5) | 20.9 | (11.3, 25.8) | > 0.05 |
| SP2027 | 26.1 | (11.6, 79.7) | 21.4 | (8.8, 29.4) | > 0.05 |
| SP2194 | 8.4 | (5.2, 35.7) | 8.0 | (5.5, 10.0) | > 0.05 |
| StkP | 2.3 | (1.0, 14.6) | 5.8 | (1.2, 63.8) | > 0.05 |
| LytB | 19.1 | (6.7, 69.3) | 16.8 | (8.1, 17.0) | > 0.05 |
| PcpA | 244.3 | (58.2, 501.5) | 242.2 | (76.6, 538.7) | > 0.05 |
| PhtD-2 | 258.3 | (120, 406.4) | 316.7 | (92.9, 479.2) | > 0.05 |
| PhtE | 1.6 | (0.0, 17.1) | 3.4 | (0, 15.5) | > 0.05 |
| PiuA | 2.8 | (1.2, 16.1) | 2.5 | (1.6, 5.4) | > 0.05 |
| PspA F2 | 0.8 | (0.3, 2.2) | 0.9 | (0.6, 2.0) | > 0.05 |
| RrgA-T4^b^ | 14.6 | (7.1, 41.0) | 11.5 | (8.9, 13.5) | > 0.05 |
| RrgB-6B^b^ | 10.9 | (5.1, 42.0) | 9.4 | (6.8, 10.9) | > 0.05 |
| Ply-1^c^ | 11.5 | (5.9, 38) | 10.6 | (9.1, 14.7) | > 0.05 |
| NanA | 48.7 | (19.7, 95.3) | 26.8 | (9.2, 59.0) | > 0.05 |
| PiaA | 7.7 | (2.7, 15.3) | 73.8 | (28.0, 153.5) | **<0.0001** |
| Ply-2^c^ | 2.4 | (1.5, 6.4) | 4.1 | (2.6, 7.5) | > 0.05 |
| PspA F1 | 8.8 | (0.8, 534.7) | 3037 | (813.4, 6848) | **<0.0001** |
| RrgB-23F^b^ | 16.8 | (7.6, 52.3) | 29.7 | (16.8, 39.3) | > 0.05 |
| RrgB-T4^b^ | 9.6 | (5.2, 30.5) | 8.6 | (7.0, 10.8) | > 0.05 |
| Spr0057 | 52.5 | (5.1, 100.4) | 16.3 | (9.0, 104.7) | > 0.05 |
| Spr0096 | 7.4 | (3.1, 21.7) | 7.4 | (5.3, 7.9) | > 0.05 |
| Spr2021 | 0 | (0, 0.4) | 0 | (0, 0) | > 0.05 |

^a^Comparison of Pnc + Adjuvant vs Pnc + WCV, Mann-Whitney test. Significant findings (p < 0.05) are shown in bold.

^b^These antigens are subunits of pneumococcal pilus-1, which is not present in either the WCV or pneumococcal EF3030 strain used in this study

^c^Ply-1 and Ply-2 are different variants of pneumolysin
